# Non-random associations of maternally transmitted symbionts in insects: The roles of drift versus co-transmission and selection

**DOI:** 10.1101/364653

**Authors:** Mathé-Hubert Hugo, Heidi Kaech, Corinne Hertaeg, Christoph Vorburger

## Abstract

Virtually all higher organisms form holobionts with associated microbiota. To understand the biology of holobionts we need to know how species assemble and interact. Controlled experiments are suited to study interactions between particular symbionts, but they can only inform about a tiny portion of the diversity within each species. Alternatively, interactions can be inferred from associations among symbionts in the field that are more or less frequent than expected under random assortment. However, random assortment may not be a valid null hypothesis for maternally transmitted symbionts in finite populations, where drift alone can result in associations. Here we report results from a European field survey of endosymbionts in the pea aphid (*Acyrthosiphon pisum*), and we develop a model to study the effect of drift on symbiont associations under different population sizes, considering varying rates of horizontal and maternal transmission. The model showed that even though horizontal transmissions and maternal transmission failures tend to randomise symbiont associations, drift can induce significant departures from random assortment, at least in moderate-sized populations. Based on these results, we carefully interpret our field survey and we re-visit the association between *Spiroplasma* and *Wolbachia* in *Drosophila neotestacea* reported by Jaenike *et al*. (2010). For this and for several significant associations between symbionts in European pea aphids we conclude that under reasonable assumptions of effective population size, they are indeed likely to be maintained by biased co-transmission or selection. Our study shows that formulating appropriate null expectations can strengthen the biological inference from co-occurrence patterns in the field.

## Introduction

Some of the interactions between organisms are so tight and durable that a new level of organisation has been defined to describe them: the holobiont (Queller & Strassmann 2016). These interactions are rarely bipartite and instead generally involve a host with a microbial community of varying degree of complexity. From the host’s perspective, these associations often lead to the acquisition of novel traits, allowing the host to expand its ecological niche (e.g., Oliver *et al*. 2010; Henry *et al*. 2013; Brucker & Bordenstein 2012). Understanding the evolutionary ecology of these interactions requires identifying how species assemble to form holobionts, both at the ontogenetic and evolutionary levels.

Large-scale screens for well-known species like *Wolbachia*, *Cardinium* or *Spiroplasma* suggest that the majority of arthropod species are infected with heritable endosymbionts (Zchori-Fein & Perlman 2004; Duron *et al*. 2008; Hilgenboecker *et al*. 2008; Regassa 2014). However, there is considerable variability in the effects these symbionts have on their hosts and in their prevalence among species. *Wolbachia* is probably the most widespread of these endosymbionts. It has been estimated to occur in 66% of arthropod species, and it typically has either low (<10%) or very high (>90%) prevalence within species (Hilgenboecker *et al*. 2008). *Wolbachia* is mainly known as a reproductive parasite (Werren *et al*. 2008), but it may also protect its host against parasites (e.g., Hedges *et al*. 2008; Teixeira *et al*. 2008; Faria *et al*. 2016) and is sometimes necessary for successful offspring production (Dedeine *et al*. 2001; Kremer *et al*. 2009). Other widespread endosymbionts of arthropods are bacteria of the genus *Spiroplasma*, infecting 4-7% of species, often with a low prevalence (Duron *et al*. 2008; Regassa 2014). Known effects of *Spiroplasma* also include reproductive parasitism (e.g.: Tabata *et al*. 2011; Sanada-Morimura *et al*. 2013; Anbutsu *et al*. 2016) as well as defense against at least three different kinds of parasites (Lukasik *et al*. 2013; Xie *et al*. 2014; Ballinger & Perlman 2017; Frago *et al*. 2017).

The pea aphid, *Acyrthosiphon pisum*, is one of the main biological models of endosymbiosis. It can be host to at least eight facultative heritable endosymbionts (Vorburger 2018), including *Spiroplasma*. Interestingly, Ferrari *et al*. (2012) showed that the communities of facultative symbionts differed strongly among host plant-associated biotypes of the pea aphid (Peccoud *et al*. 2009), although the prevalence of *Spiroplasma* is only weakly affected by biotype, which explains only 9% of the variance (Ferrari *et al*. 2012). A symbiont can spread in a host population as a reproductive parasite, for which there is some limited evidence from pea aphid *Spiroplasma* (Simon et al. 2011), or by providing a benefit to offset the cost it inflicts on the host. For example, *Spiroplasma* may protect pea aphids against entomopathogenic fungi (Łukasik *et al*. 2013) or parasitoid wasps (Frago *et al*. 2017). However, this cost-benefit balance varies depending on the environment, which is thought to be the main reason for the observed polymorphism of facultative symbiont communities. For example, defensive symbioses depend on the presence of some parasites of the host, and some symbioses help the host to cope with warm environments (e.g., Russell & Moran 2006). The cost-benefit balance may also depend on the associations with other symbionts. If two symbionts provide the same service, then one of them might be redundant an thus too costly to the host. This may be the reason why defensive bacterial symbionts are less frequent in aphids protected by ants (Henry *et al*. 2015), or why the two defensive symbionts *Serratia symbiotica* and *Hamiltonella defensa* rarely co-occur in pea aphids (Oliver *et al*. 2006). Also, interactions between symbionts can lead to non-additive effects in a symbiont- and host-specific manner, making the outcome of a given association difficult to predict. For instance, in *A*. *pisum*, *H*. *defensa* increases the titer of *S*. *symbiotica*, but *S*. *symbiotica* does not affect the titer of *H*. *defensa* (Oliver *et al*. 2006). In the presence of *Spiroplasma*, *H*. *defensa* decreases the fecundity of its host *A*. *pisum* while it increases the fecundity of the aphid *Sitobion avenae* (Lukasik *et al*. 2013).

Interactive effects that vary from one symbiont strain to the other limit the utility of controlled laboratory experiments, which usually include only a few particular strains, for making predictions about the overall interactions among symbionts in natural populations. For this reason, results from controlled experiments are often compared to analyses of field surveys (for several examples, see Zytynska & Weisser 2016). These analyses notably aim at identifying pairs of symbionts for which the co-occurrence is more or less frequent than expected under the null hypothesis of random assortment (hereafter, positive and negative associations). Three kinds of mechanisms are generally considered when trying to explain such deviation from random assortment. Firstly, the symbionts could interact in a way that increases or decreases the rate of maternal transmission failures (e.g. Rock *et al*. 2017), which should lead to negative or positive associations, respectively. Secondly, the symbionts could have an interactive effect on host fitness, enhancing or hindering their co-transmission to the next generation (e.g. Oliver *et al*. 2006). Thirdly, Jaenike (2012) and Smith *et al*. (2015) suggested that neutral or even slightly costly maternally transmitted symbionts could spread in the host population by chance association with another symbiont that is beneficial to the host. Faithful maternal transmission would maintain the association between the two symbionts even if it were advantageous for the host to lose one. This symbiont hitchhiking is analogous to the genetic hitchhiking (or draft), where a neutral or slightly deleterious mutation spreads in the population because of its linkage disequilibrium with a beneficial mutation (Felsenstein 1974). This symbiont hitchhiking might be responsible for the evolutionary maintenance of the symbiont called *X*-*type*, which is costly to its host, has not been found to provide any counterbalancing benefit, but is positively associated with the defensive symbiont *H*. *defensa* (Doremus & Oliver 2017).

Jaenike (2012) argued that because most symbionts show some degree of maternal transmission failure, associations due to symbiont hitchhiking should disappear rapidly. Thus, in most cases, the presence of positive (or negative) associations between symbionts suggests an interaction that favors (or hinders) their co-occurrence. Jaenike *et al*. (2010) showed that *Spiroplasma* and *Wolbachia* in *Drosophila neotestacea* are positively associated despite imperfect maternal transmission. By combining these observations with a mathematical model, they suggested that these two symbionts are likely to be interacting positively with each other. As we will show in this paper, positive and negative associations are also expected to appear and persist by drift, implying that without information about the effective female population size, one needs to be cautious in assigning biological meaning to such associations.

In the first part of this study, we used a field survey of *A*. *pisum* symbiotic infections to identify positive and negative associations among symbionts. We report some new associations and confirm some already known associations. We also show that there are three sub-clades of *Spiroplasma* in pea aphids, which tend to be associated with very different symbiont communities. In the second part of this study, we developed a simulation model of maternally and horizontally transmitted symbiont communities in female host populations of finite size. We used this model to assess the effect of drift on the tendency of symbionts to be randomly assorted to each other or not, thus aiding our interpretation of observed associations in the field. We also applied the model to the association between *Spiroplasma* and *Wolbachia* in *D*. *neotestacea* described by Jaenike *et al*. (2010) as a useful test case, and, under a reasonable assumption for its effective population size, we come to the conclusion that this association is indeed likely to reflect a positive interaction between symbionts.

## Materials and Methods

Natural symbiont co-occurrence

### Field sampling and symbiont screening

We sampled 498 aphids in France, Switzerland, Germany and Denmark during autumn 2014 and spring and summer 2015. We selected colonies that were at least 2 meters apart from each other to limit the risk of sampling offspring of the same mother. For each sample, we recorded the host plant and the GPS coordinates. We characterised the presence of seven known facultative endosymbionts by diagnostic PCR using symbiont-specific primers to amplify a part of the 16S rRNA gene (Table S1). DNA was extracted from individual aphids using the ‘salting out’ protocol (Sunnucks & Hales 1996) and the PCR cycling conditions are described by Henry *et al*. (2013). We also ran a diagnostic PCR for the obligate endosymbiont *Buchnera aphidicola*, which is present in all aphids and thus served as an internal positive control for the quality of the DNA preparation. Only samples testing positive for *B*. *aphidicola* were included in the final dataset.

### Phylogeny of *Spiroplasma* in pea aphids

Because we had a special interest in *Spiroplasma* infecting pea aphids (Mathé-Hubert et *al*. in prep.), we also analysed the distribution of intraspecific diversity in this symbiont. This analysis included the 26 strains found in the above-mentioned field sampling as well as 11 strains that were kindly provided by Ailsa McLean (Department of Zoology, University of Oxford, UK; Table S2). The diversity of *Spiroplasma* was characterised using a phylogeny based on the dnaA gene and the rpoB gene, including its surrounding regions (445 pb and 2731 pb, respectively, including primers; Table S1). The PCR cycling conditions are as described by Henry *et al*. (2013), except that the elongation time of the primer pair RpoBF1.ixod was doubled (Table S1). We deposited all dnaA and rpoB sequences in Genbank (accession numbers: MG288511 to MG288588).

We inferred the phylogenetic tree of the 37 *Spiroplasma* strains using *Spiroplasma sp*. in *Ostrinia zaguliaevi* as outgroup, which is the most closely related species to *Spiroplasma* of the pea aphid for which we could obtain sequences for both the rpoB and dnaA genes. The substitution model “GTR + gamma + invariant sites” was identified by AICc (“phangorn” R package v 2.2.0, Schliep 2011) as the best-fitting for the MAFFT-aligned and then concatenated dnaA and rpoB sequences. It was used to build a maximum likelihood tree with the software MEGA6 (Tamura *et al*. 2013). This analysis identified three main clades of *Spiroplasma* from pea aphids that are later referred to as clade 1, 2 and 3.

### Statistical analysis

All analyses were performed using the R software (version 3.4.4; R Core Team 2018). To detect associations of symbionts that are more or less abundant than expected under random assortment while accounting for spatiotemporal non-independence among samples we predicted the presence or absence of each symbiont species with a regression random forest model (RF). In each RF, the following explanatory variables were used: latitude, longitude, season (number of days since the start of the year), host plant on which the aphid has been sampled, aphid colour (pink or green), presence or absence of the six other symbionts (one variable per symbiont) and the total number of other symbiont species infecting the aphid.

In order to handle the multicollinearity as accurately as possible, RFs were grown with conditional inference trees and the effect of variables estimated with conditional importance (package *party*; Hothorn *et al*. 2006). Our methodology mostly follows the advice of Jones and Linder (2015). The only difference is that we applied the permutation approach developed by Hapfelmeier & Ulm (2013) to the conditional importance of explanatory variables to estimate their *p*-values that were then adjusted to keep the false discovery rate at 5% (Benjamini & Yekutieli 2001). With this approach, there is one model per symbiont, the presence or absence of the focal symbiont being explained by the presence or absence of the six other symbionts. Thus each pair of symbiont (a and b) is considered by two models, one explaining the symbiont a and the other explaining the symbiont b. This approach thus leads to two *p*-values per couple of symbionts. To facilitate the interpretations of the results, we repeated this analysis by restricting the dataset to aphids sampled on *Medicago sativa* and to aphids sampled on *Trifolium spp*., which represent 30 and 33% of all field samples, respectively. This restriction avoids lumping together aphids from multiple host races and thus simplifies the interpretation. We refer hereafter to these three types of models as RF_WD_ (whole dataset), RF_M_ (*M*. *sativa*) and RF_T_ (*Trifolium spp*.). For RF_M_ and RF_T_ the host plant was removed from the set of explanatory variables.

These analyses revealed that some symbionts are less frequent in aphids already containing other symbiont species, while others are not significantly affected by the presence of other symbionts. To further investigate this, we tested if there was a link between the frequency of each symbiont species and the average number of additional symbiont species with which it co-occurs. Such a link is expected to occur even in the absence of interaction between symbionts, because under occasional horizontal transmission, common symbionts are more likely to become associated with their own kind than rare symbionts. We used a Wald test (function *regTermTest* of the package *survey*; Lumley & Scott 2014) to test if the observed slope of the relationship between symbiont frequency and the mean number of co-infecting symbionts was different from the slope expected under the null assumption of no interaction between symbionts. The formula used to compute the expected slope is provided in Appendix S1.

To investigate the intraspecific distribution of *Spiroplasma*, we used a classification RF to predict the phylogenetic clade of *Spiroplasma* for each *Spiroplasma* infected aphid. The *Spiroplasma* strain S362 was excluded from the analysis because it was not assignable to one of the three phylogenetic clades. The variables were the same as in the previous analysis except that the aphid colour was not available for all aphids and was thus not used. For significant variables, we compare clades to each other in a pairwise fashion to characterise inter-clade differences. Since some of these significant variables are continuous and some are categorical, we used Wilcoxon or Fisher’s exact tests, respectively to perform these pairwise comparisons.

### Simulations of the symbiont co-occurrences

We investigated the effect of drift on deviations from random assortment of symbionts by simulating populations of hosts with two species of maternally transmitted symbionts. In short, we simulated populations of female hosts reproducing with non-overlapping generations, and being infected by zero, one or two symbionts species (or strains). Symbionts are maternally transmitted with a varying degree of efficiency and they are also horizontally transmitted at varying rates. Specifically, we simulated 3000 replicates of all combinations of the following sets of parameters: Effective female population sizes (N_e_F: 10^3^, 10^4^, 10^5^, 10^6^, and 10^7^), Successful maternal transmission rates (M_T_: 1, 0.999, 0.99, 0.90), Horizontal transmission rates (i.e.: Average number of horizontal transmission events caused by each infected host; H_T_: 0, 0.001, 0.01, 0.1).

To initiate the populations we first randomly chose the frequencies of the two symbiont species from a uniform distribution and computed the frequency of the four kinds of symbiont communities (no symbiont, symbiont 1, symbiont 2, symbionts 1 and 2) according to the assumption of random assortment and the frequencies of the symbionts. Then, the population sizes of the four kinds of communities were sampled in a multinomial distribution, according to the previously computed probabilities.

Population evolution was simulated for 10^5^ generations or stopped if the polymorphism of infection was lost. In the absence of selection, for most combinations of M_T_ and H_T_, the symbionts get rapidly fixed or lost which prevents assessing deviations from random assortment. To limit this loss of polymorphism, we have set the direction and the strength of the selection on the presence of each symbiont in a way that the expected equilibrium frequency of each symbiont equals their initial frequency. For more detail, see Appendix S2. This procedure was more efficient at delaying the loss of polymorphism due to the maternal transmission failure, than the loss due to horizontal transmissions. This is because the amount of horizontal transmissions has a positive feedback on itself, while the amount of vertical transmission failure has a negative feedback on itself (Appendix S2).

At each generation, 500 individuals were randomly sampled from the population and used to test the significance of the deviation from the assumption of random assortment using a χ^2^-test (Yates 1934) and to assess the sign of the deviation. The *p*-values are recorded at generations 0, 10, 10^2^, 10^3^, 10^4^ and 10^5^. We also assessed if, as it is often assumed, associations lasting for several years are unlikely driven by drift. We estimated the number of generations needed for a previously significantly positive association to become significantly negative. This was computed for each replicate as the number of simulated generations divided by the number of inversion of the sign of significant trends. The detailed description of the model and the model itself are in Appendix S2.

Jaenike *et al*. (2010) argued that *Wolbachia* and *Spiroplasma* in *D*. *neotestacea* are probably interacting in a way that enhances the fitness of co-infected hosts because they are positively associated in natural populations, despite having a maternal transmission rate of around 0.96, which should rapidly randomise them. We used our model to assess how robust this conclusion is to drift. We simulated, as described above, populations of female hosts infected by up to two symbionts with initial frequencies of 0.8135 and 0.4289, which are the mean frequencies of infection with *Wolbachia* and *Spiroplasma* in *D*. *neotestacea* measured by Jaenike *et al*. (2010). In this simulation, we set the maternal transmission rate to 0.96, which is the mean of the estimates obtained by Jaenike *et al*. (2010) for both *Wolbachia* and *Spiroplasma*.

## Results

### Natural co-occurrence of pea aphid symbionts

The RFs analysing the associations of symbionts in all field-sampled aphids revealed three positive associations and six negative associations. Of these associations, all were detected in RF_WD_ (Fig. 1A) six were detected in RF_T_ (Fig. 1B), and only two were detected in RF_M_ (Fig. 1C). The lower number of significant associations in *M*. *sativa* (RF_M_) than in *Trifolium spp*. (RF_M_) is unlikely due to a lower statistical power since the sample sizes and the average number of symbionts per aphid were similar in these two groups (*M*. *sativa*: 148 aphids with 0.97 symbionts per aphid on average; *Trifolium*: 161, 0.77). Of the 11 significant associations already identified by other studies on pea aphids, six were also found in this study, and all associations reported by several studies (including ours) were always of the same sign (Table 1).

**Figure 1:**
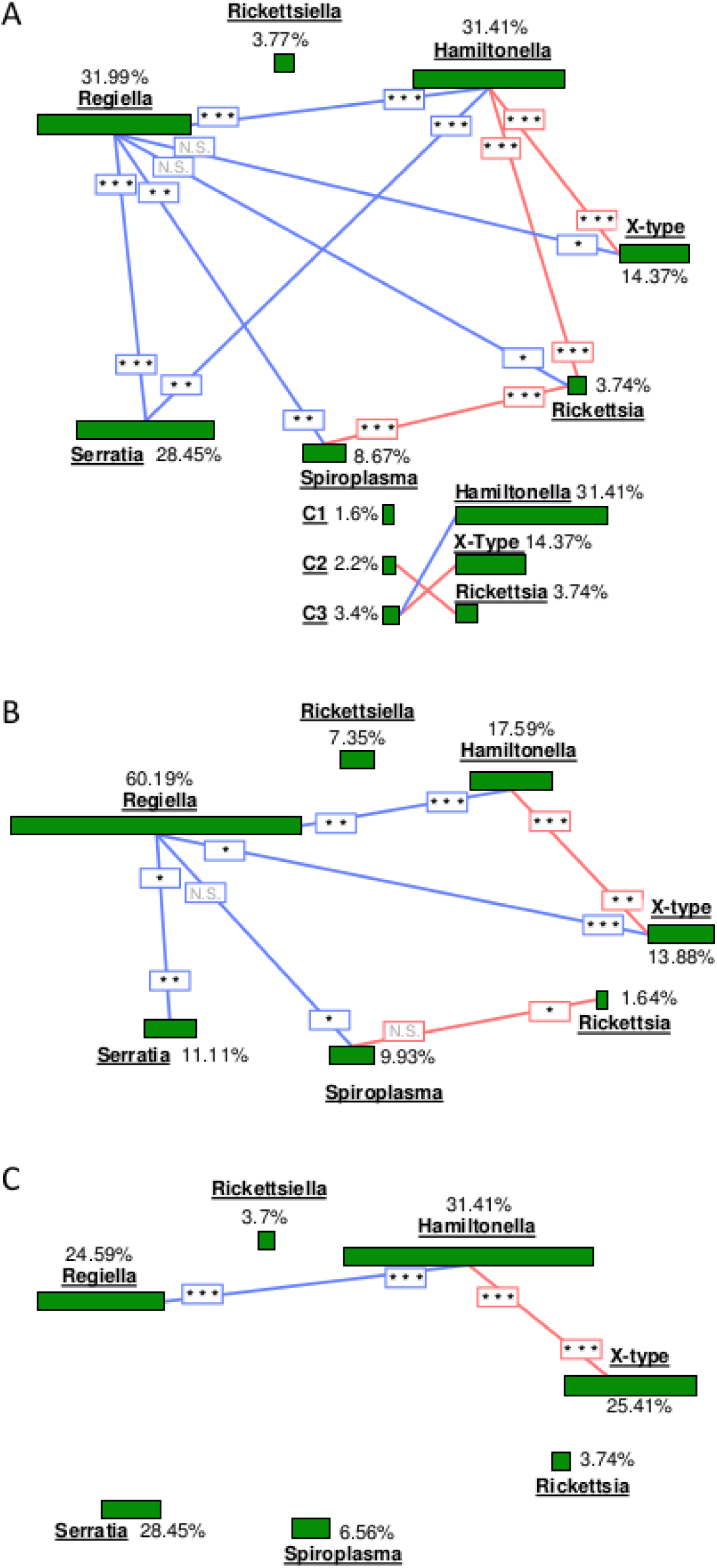
Patterns of symbiont co-occurrence. The seven symbiont species are represented by green boxes whose size is proportional to the overall prevalence of the symbiont in the whole dataset (A; N=498), in aphids from *Trifolium spp*. (B; N=161) and in aphids from *Medicago sativa* (C; N=148). Red and blue lines connect symbionts that co-occur more or less often than expected under random assortment, respectively. Stars indicate the FDR-adjusted level of significance of these associations and are placed close to the symbiont that was the dependent variable in the random forest models.

**Table 1:**
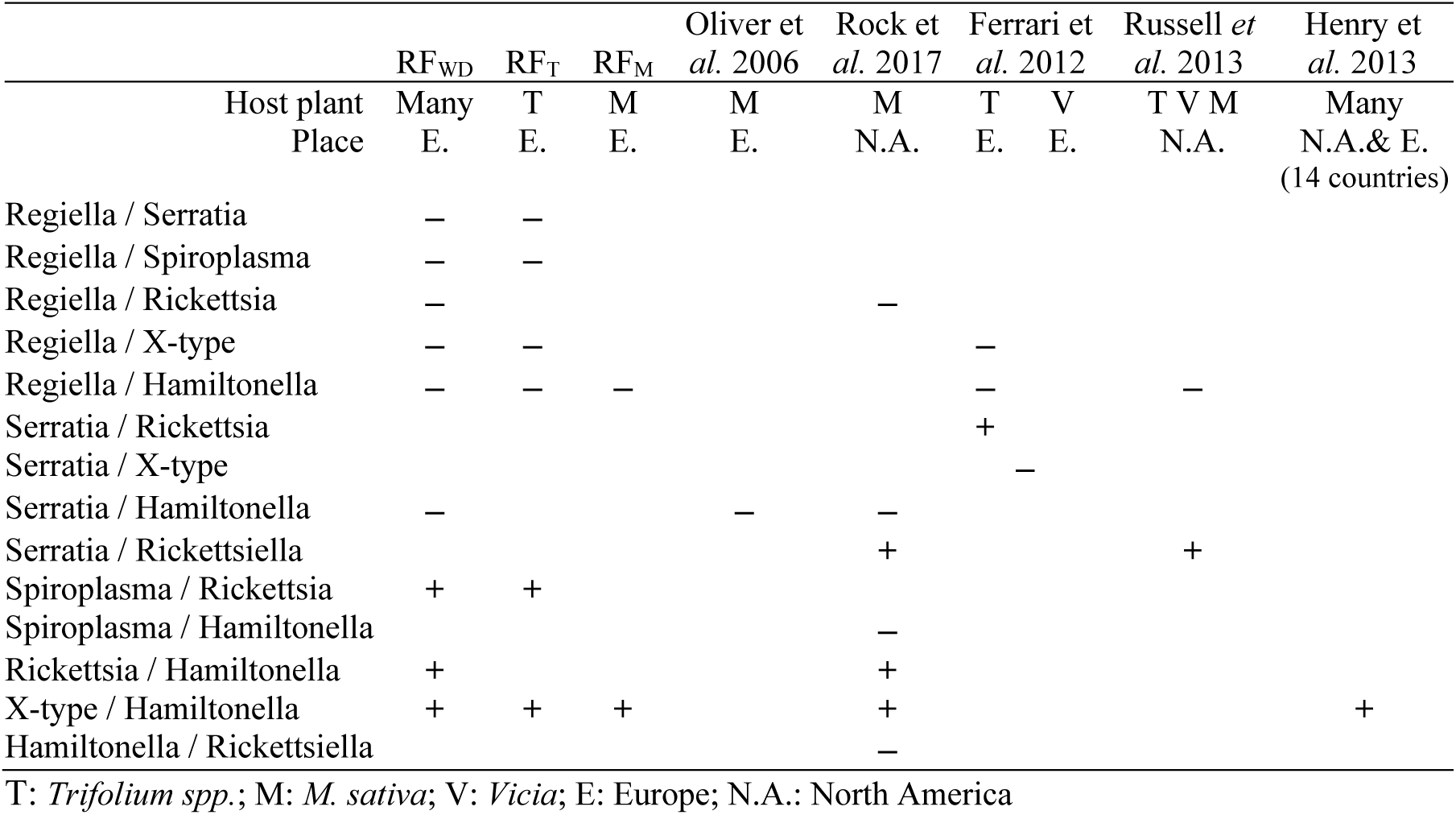
Patterns of symbiont co-occurrence in this study and in other studies on pea aphids.

Some symbiont prevalences co-varied negatively with the total number of co-infecting symbiont species, whatever their identity (*H*. *defensa*: FDR-adjusted *p*-values: *P*=0.002 and P=0.02 in RF_WD_ and RF_M_, respectively; *R*. *insecticola*: *P*<0.001 in the three models RF_WD_, RF_T_ and RF_M_; *S*. *symbiotica*: *P*<0.001 and in both RF_WD_ and RF_M_). For pea aphids from *Trifolium spp*., the relationship between symbiont prevalence and the mean number of co-infecting symbionts was tight (*R*^2^=0.98) and its slope was more negative than the expected slope under the assumption of no interactions among symbionts (Wald test: *P*<0.001, Fig. 2B), but this was mostly driven by *R*. *insecticola*. However, repeating the analysis without aphids infected by *R*. *insecticola* also yielded significantly different slopes (*P*=0.04; R^2^=0.88), mainly because *Rickettsiella* and *Rickettsia*, had a low frequency and tended to occur in aphids already infected by more than one other symbiont species (Fig. 2C). For pea aphids from *M*. *sativa*, the slope was also more negative than expected, but the difference was not significant (*P*=0.45, Fig. 2A).

**Figure 2:**
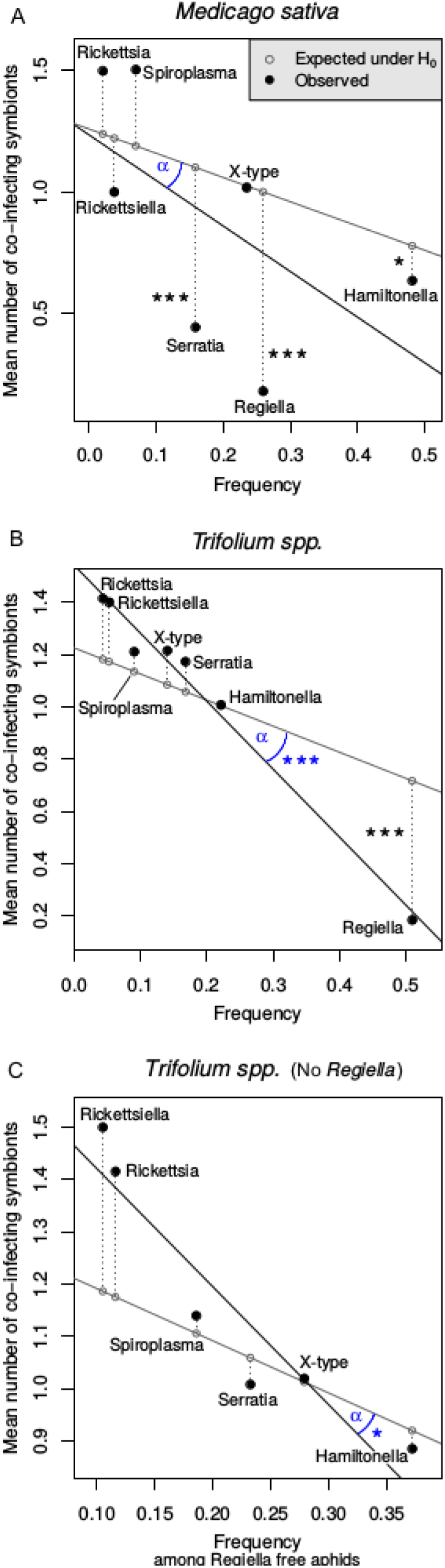
Relationship between symbiont frequency and mean number of other symbionts species. Comparison of the actual (black) and expected (grey) relationship between the frequency of endosymbionts species and the mean number of other symbiont species with which they co-occur. Each observed value is connected to its expected value by a dotted line. Stars along these lines indicate the level of significance detected by RF models. Analysis were performed on pea aphids from *Medicago sativa* (A) and *Trifolium spp*. (B). Panel C refers to the analysis performed on aphids from *Trifolium spp*., but excluding individual infected with *Regiella insecticola* from the analysis. For each of these three cases, we tested if the angle between the two slopes (α) differed significantly from zero.

All these models included the variables longitude, latitude, season and in the case of RFWD, the host plant, to account for the non-independence between samples. However, these variables are highly correlated, and although we used conditional inference trees and conditional importance, the results should be interpreted with caution. The effects of these four variables on the frequency of each symbiont are described in Figure S1.

### *Spiroplasma* intraspecific diversity

The pairwise comparison of sequences from *Spiroplasma* of pea aphids revealed that on average 0.63% of sites were different (max divergence=1.2%) which corresponds to 38% of the mean divergence between *Spiroplasma* of pea aphids and *Spiroplasma* of the moth *Ostrinia zaguliaevi* (1.6% of sites being divergent). The phylogenetic tree suggests that in Europe, *Spiroplasma* of the pea aphid is sub-divided into at least three clades, but clade 3 in particular has a low bootstrap support (Fig. 3). The relative frequencies of these three clades did not depend on the host plant (non-FDR-adjusted *P*=0.98; Fig. 3) but were strongly dependent on the symbiont community. Clade 2 was more frequent in aphids already infected by other endosymbionts (FDR-adjusted *P*=0.01) than the other two clades. This was marginally non-significant when comparing it to cade 1 and marginally significant when comparing it to clade 3 (*P*=0.06 and *P*=0.03, respectively; Wilcoxon-test). The *Spiroplasma* clades were also differently associated with *H*. *defensa*, *X*-*type* and *Rickettsia* (FDR-adjusted *P*=0.02, 0.003 and 0.003, respectively). Specifically, clade 3 co-occurs less frequently with *H*. *defensa* than clades 1 and 2 (*P*=0.02 and 0.01; Fisher-exact test) and more frequently with *X*-*type* than clades 1 and 2 (*P*=0.003 and 0.006; Fisher-exact test; Fig. 1 and 3). Also, clade 2 is more frequently associated with *Rickettsia* than clades 1 and 3 (*P*<0.001 and <0.001; Fisher-exact tests; Fig. 1 and 3).

**Figure 3:**
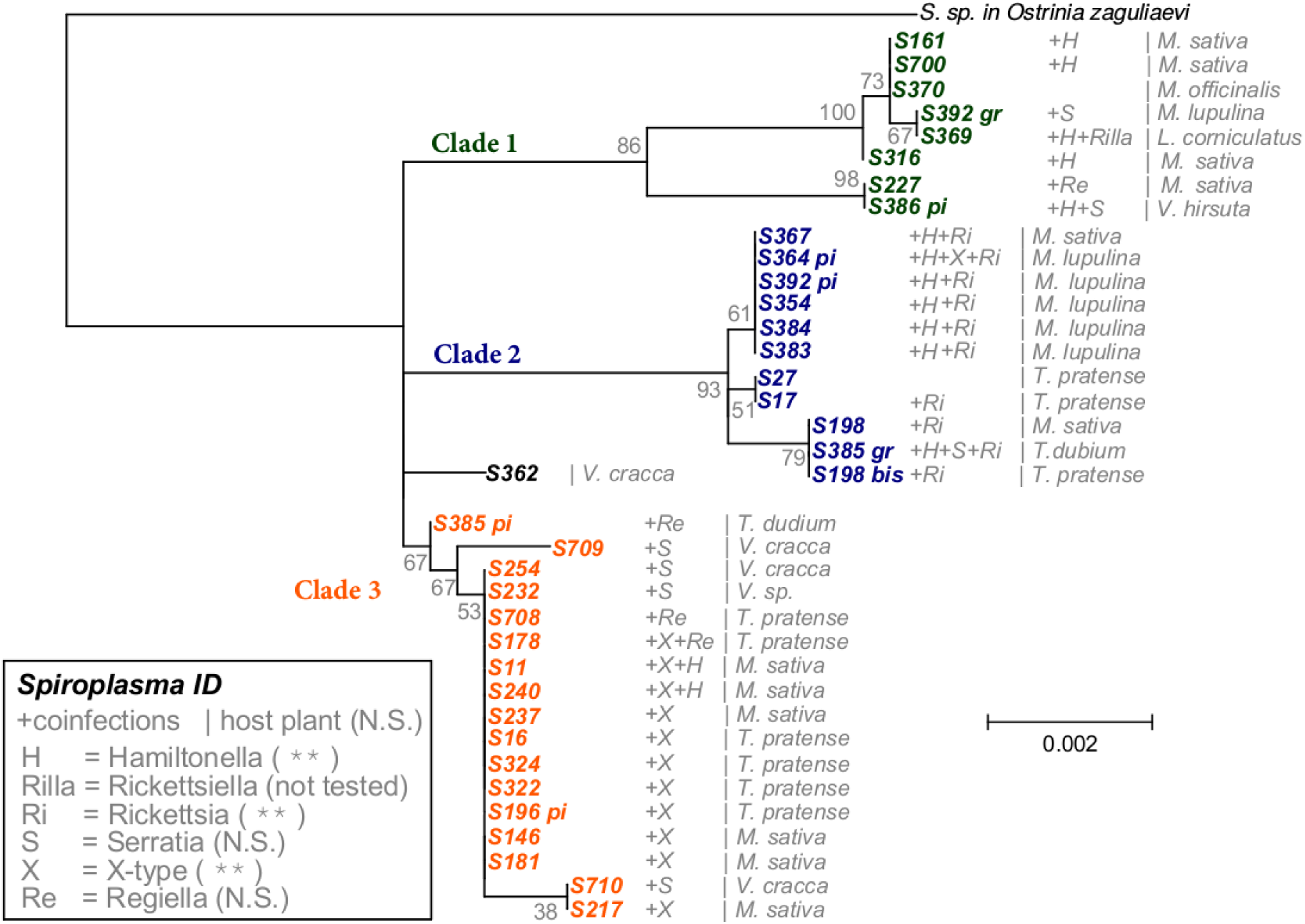
Phylogenetic tree of *Spiroplasma* from pea aphids. Maximum likelihood phylogenetic tree of *Spiroplasma* strains from pea aphids, reconstructed from concatenated rpoB and dnaA sequqences and using *Spiroplasma* from *Ostrinia zaguliaevi* as outgroup. Values in grey are the bootstrap support for the tree topology. The lists of the co-infecting symbionts and the host plants are indicated on the right side of the strain name. Scale bar indicates the substitution rate. The legend gives the abbreviations of the symbiont names, as well the FDR-adjusted level of significance of these variables in the RF predicting the clade of each *Spiroplasma* strain (1, 2 or 3).

### Simulations of the symbiont co-occurrences evolving by drift

Symbiont associations that are more or less frequent than expected under the hypothesis of random assortment are generally interpreted as a sign that an interaction between the symbionts promotes or prevents their co-occurrence. Our simulations showed that for symbionts with perfect maternal and no horizontal transmission, drift always leads to strong deviations from random assortment (Fig. 4). Such deviations take a longer time to appear in large populations where drift is weak. Also, the median number of generations between the appearance of a significant deviation from random assortment of a given sign and the inversion of the sign is 54, 117, and 211, respectively, for effective female population sizes of 10^3^, 10^4^ and 10^5^ or more (Fig. S4). As often presumed, less-than-perfect maternal transmission and horizontal transmissions tend to randomize symbiont associations (Fig. 4 and S3).

**Figure 4:**
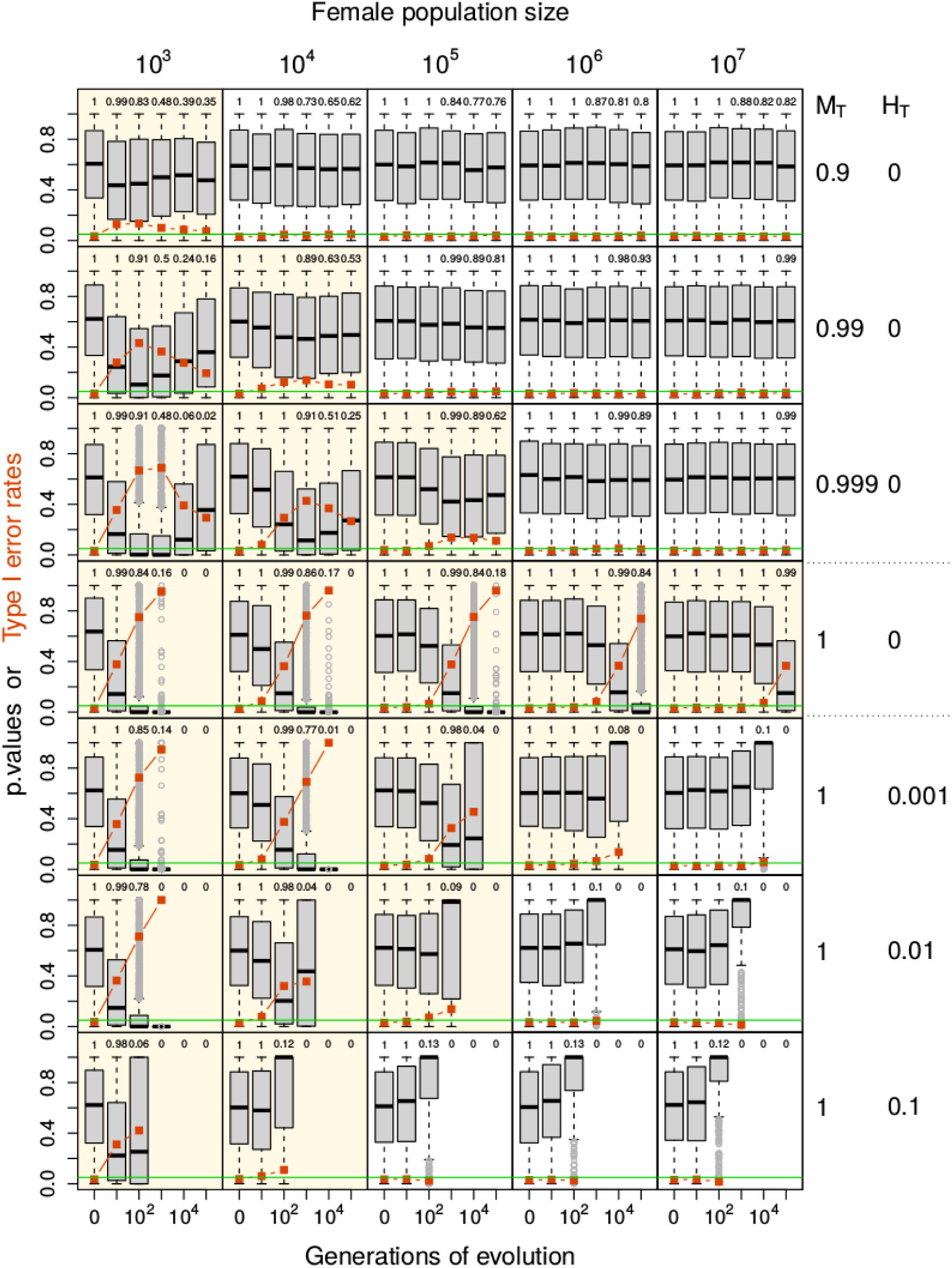
Deviations from random assortment induced by drift. The frequency of two maternally transmitted symbionts evolved for up to 10^5^ generations, starting from a population in which symbionts were randomly assorted. Boxplots show the *p*-values of χ^2^-tests assessing the deviations random assortment at generations 0, 10, 10^2^, 10^3^, 10^4^ and 10^5^. Each set of boxplots corresponds to 3000 populations evolving with the combination of the parameters indicated aside: ‘female population size’ (columns), ‘horizontal transmission rate’ (H_T_; rows) and ‘maternal transmission rate’ (M_T_; rows). The green horizontal line shows the 0.05 threshold, and the orange squares and lines indicate the type 1 error rate. Analyses of field surveys testing for deviation from random assortment usually assume that the type 1 error rate is 0.05. Combinations of parameters where this is not the case have a yellowish background. The numbers above the boxplots indicate the proportion of populations that still retained some polymorphism of infection by both symbionts.

However, our model shows that this effect can be offset by drift. Thus, whether random assortment is a valid expectation for non-interacting symbionts will depend on the effective female population size as well as on the rates of horizontal transmissions and maternal transmission failures (Fig. 4 and S3). Horizontal transmission and maternal transmission failure are also shortening the mean number of generations between inversions of the sign of deviations from random assortment (Fig S3). In case of a maternal transmission rate of 0.9, and a horizontal transmission rate of 0.1, the median number of generations is 38, 50, and 52, respectively, for effective female population sizes of 10^3^, 10^4^ and 10^5^ or more.

Our simulations suggest that the observations reported by Jaenike *et al*. (2010) would be compatible with drift if the effective female population size of *D*. *neotestacea* would be 10^3^, but not if it would be 10^4^ or more. Indeed, for the symbiont frequency and maternal transmission rate reported by Jaenike *et al*. (2010) and a female population size of 10^3^, the type 1 error rate of a test of independence of symbiont occurrence is 14%, while it is around 5% for female population size of 10^4^ or more (Fig. S4). Our simulations also suggest that on average, the sign of deviations from random assortment that are driven by drift would only happen every 34, 48, and 52 generations, respectively, for female populations size of 10^3^, 10^4^, and ≥ 10^5^ (Fig. S4).

## Discussion

Understanding how symbionts associate and interact within a host is important but challenging. Laboratory experiments address this question by controlling all relevant parameters and observing the outcomes, but they can only accommodate a tiny portion of the natural diversity of each interacting species. In addition, such studies have often found that the outcome depends on the genotypes of the interacting partners (e.g.: Russell & Moran 2006; Oliver *et al*. 2009; Vorburger & Gouskov 2011; Hansen *et al*. 2012; Lukasik *et al*. 2013; Weldon *et al*. 2013; Niepoth *et al*. 2018), further complicating predictions about these interactions in natural populations. Comparisons with field observations are therefore essential. When analysing field surveys, interactions between symbionts are tentatively inferred by comparing the observed frequency of co-occurrences to the frequency expected under the hypothesis of random assortment. Departures from random assortment have been reported frequently (e.g., Ferrari *et al*. 2012; Henry *et al*. 2013 in addition to the above-mentioned studies). Of the 21 possible pairwise associations among the seven facultative endosymbionts considered here, 11 have already been reported to have significantly higher or lower frequencies than expected under random assortment in earlier studies on pea aphids (Table 1). Six of these associations were also found in our field sampling, and three are reported for the first time. When focusing on *Spiroplasma*, we even found significant associations at the intra-specific level. The three main *Spiroplasma* clades identified in the phylogenetic tree were non-randomly associated with other symbionts, independent of the host plants the aphids were collected from. Such intraspecific variation in a symbiont-symbiont association has also been reported between *X*-*type* and *H*. *defensa* in the pea aphid (Doremus & Oliver 2017). But what is the biological meaning of these associations?

### Drift induces deviations from random assortment

Our simulation model showed that random drift also induces associations among maternally transmitted symbionts, suggesting that random assortment is not an appropriate null model to compare symbiont co-infections against. The reason for that is most easily understood by considering the coalescence framework. Statistical tests used to detect departures from random assortment assume that samples are independent of each other. While this may apply to horizontally transmitted symbionts, it will not apply to maternally transmitted symbionts. Some individuals will have the same symbiont simply because they share a female ancestor that transmitted this particular symbiont community to all of its offspring. In population genetics, this phenomenon is referred to as the coalescence (Balding *et al*. 2007; which should not be confounded with the ‘community coalescence’; Rillig *et al*. 2015). One of the measures of the strength of drift is the expected coalescent time, the average number of generations between two randomly sampled alleles and their most recent common ancestor. It is equal to 2N_e_ for diploid autosomal genes, but it is only N_e_/2 for maternally transmitted cytoplasmic genomes (assuming a sex-ratio of 0.5). This is because only females do transmit the cytoplasmic genome, and they have only one copy of it (Moore 1995; Jaenike 2012). Cytoplasmic genomes, including endosymbionts, hence undergo four times more drift than nuclear autosomal genes.

Jaenike (2012) investigated how the population genetics framework can be adapted and used to study the evolution of communities of maternally transmitted symbionts by comparing each symbiont to a gene. However, given the generally high fidelity of maternal transmission and the low rate of horizontal transmission of endosymbionts, one could also compare the whole symbiont community to one gene with many alleles. Assuming no mutational bias, mutations increase allelic diversity and maintain the alleles at a similar frequency, while, drift has the opposite effect. This mutation-drift balance is largely analogous to the balance between maternal transmission failures, horizontal transmissions and drift that we studied with our model. The main difference is that while maternal transmission failure effectively acts as a directional mutation pressure, where the number of individuals mutating from one state (infected) to the other (uninfected) is proportional to the number of individual in the original state (infected), this is not true for horizontal transmission. The probability of undergoing a horizontal transmission increases with the frequency of the symbiont, which makes polymorphism less easily maintained in the presence of horizontal transmission.

Drift-induced deviations from random assortment can persist for a very long time. In a population of diploid autosomal genes, a neutral mutation that reaches fixation does so, on average, 4N_e_ generations after it appeared (Kimura & Ohta 1969), or after N_e_ generations in a haploid, maternally transmitted gene. Thus, we should expect that drift-induced deviations from random assortment of symbionts should also be somewhat stable in time. In agreement with that, our simulations of two strictly maternally transmitted symbionts show that drift-induced inversions of the sign of significant deviations from random assortment occur every 50 to 200 generations on average, depending on the effective female population size. These numbers should not be used as a general reference because significance depends on the size of the samples used to assess deviations from random assortment (500 hosts in our simulations). Departures from random assortment became less stable in presence of horizontal transmissions and maternal transmission failures.

### The association between *Wolbachia* and *Spiroplasma* in *Drosophila neotestacea* as a test case

Jaenike *et al*. (2010) studied the maintenance of the positive association between *Wolbachia* and *Spiroplasma* in *D*. *neotestacea*. They used a mathematical model to show that given the maternal transmission rate estimated at 0.96, the association should disappear very rapidly in the absence of any positive interactions between the two symbionts. While it is true that this relatively imperfect maternal transmission will push a population towards random assortment, their model only considered the frequency of the symbionts. Thus, it implicitly assumed an infinite population size and omitted drift which, as we have shown, will push populations towards non-random assortment. Using our model with the same parameter estimates, we show that a significant association between two symbionts could be induced just by drift if the effective female population size of *D*. *neotestacea* were in the range of 10^3^, but not if it were in the range of 10^4^ or larger. The effective population size of *D*. *neotestacea* is not known, but the effective population size of North American *D*. *melanogaster* has been estimated to be between 3 to 5 × 10^6^ (Garud *et al*. 2015). Unless the population structure of *D*. *neotestacea* is radically different from *D*. *melanogaster*, we can reasonably assume that its effective population size is higher than 10^4^. Therefore, a stronger mechanism than drift appears to be responsible for maintaining the positive association between *Wolbachia* and *Spiroplasma* in *D*. *neotestacea*, corroborating the conclusion of Jaenike *et al*. (2010) that positive selection acts on the combination of these two endosymbionts.

### Symbiont associations in pea aphids – selection or drift?

After emphasizing the importance of considering drift as a source of non-random assortment among symbionts, we return to the interpretation of positive and negative associations among facultative endosymbionts observed in pea aphids. Are they maintained by interactions among symbionts or just a consequence of drift? Good estimates of effective female population size would obviously help. Unfortunately, this is a tricky problem in aphids and other cyclical parthenogens. Although aphids can reach enormous population sizes, they undergo a bottleneck each winter, and clonal selection during the asexual phase of the life cycle (approx. 7-14 generations in pea aphids; Barker 2016) can be intense (e.g., Vorburger 2006), which will also reduce the effective population size. This clonal selection acts on the three components of genetic variance (additive, epistatic and of dominance), but the optimisation it induces on the non-additive variances is lost at each sexual generation, which maintains the presence of clonal selection from year to year (Lynch & Deng 1994). On the other hand, aphids are good dispersers and exhibit shallow genetic population structure over large geographic scales. For example, Ferrari *et al*. (2012) reported *F*_sT_-values ranging from 0.03 to 0.11 for pea aphid populations from the same host plants across different European countries, and Via & West (2008) reported a mean *F*_ST_ of 0.03 for North American populations of the pea aphid. Such high population connectivity should have a positive effect on effective population size. We do not know the effective population size of pea aphids, but DNA sequence-based estimates from other cyclical parthenogens, waterfleas of the genus *Daphnia*, are rather high (300′000 – 600′000; Haag *et al*. 2009). If estimates were similarly high for pea aphids, the importance of drift in generating non-random assortment of symbionts would be limited (Fig. 4).

Another important aspect to consider is the consistency of the sign of significant associations. While drift will generate associations of random and (slowly) fluctuating sign, selection is expected to consistently favour either positive or negative associations between particular pairs of facultative endosymbionts. For significant associations that were discovered in multiple studies, the sign of the association was always the same (Table 1). Finding particular combinations of symbionts consistently over- or underrepresented across different times and places suggests they are not caused by drift. For example, the European pea aphids population is thought to have colonised North America at least 200 years ago, which would represent 1600 to 3000 pea aphid generations, and there is strong genetic differentiation among pea aphids from the two continents today (Brisson *et al*. 2009). Despite this separation, the four associations that have been reported in both continents are of the same sign. However, any of these associations observed one time on each of the two continents are only two independent observations of a kind of event that may also happen by drift.

In addition to testing for deviations from random assortment, some studies have also assessed whether symbiont species tend to be differently associated with aphids that are already infected with 0, 1, 2 or more other symbiont species (e.g., Ferrari *et al*. 2012; Russell *et al*. 2013; Zchori-Fein *et al*. 2014; Rock *et al*. 2017). As these analyses are assuming that all the sampled symbiont communities are independent, they are also affected by drift. In our field survey, we found that *H*. *defensa*, *S*. *symbiotica* and *R*. *insecticola* occurred more frequently in aphids containing no or few other symbiont species than expected under the assumption of random assortment, although this was only significant in aphids sampled from *M*.*sativa*. When further investigating this by characterising the link between the frequency of each symbiont species and the number of co-infecting symbiont species (see Figure 2), we found that this link was non-significant absent in aphids sampled on *M*. *sativa*, and stronger than expected under the assumption of no symbiont interaction in aphids sampled on *Trifolium spp*. At least two non-exclusive mechanisms could have yielded such a pattern. Firstly, rare symbionts might be rare because they need the presence of other symbionts to persist in their host. Secondly, symbionts might have adapted to co-occurrence patterns that are largely a function of their relative frequencies, which entail that frequent symbionts should be less likely to share a host with other symbiont species than rare symbionts. Rare symbionts are thus more strongly selected for persistence in the presence of other symbiont species. This highlights that only abundant symbiont associations are efficiently optimised by natural selection. It is therefore worth considering that associations between symbionts that are currently maintained by a positive interaction may have evolved as a consequence of an association that had originally appeared by drift or draft.

Lastly, inference on the biology of particular symbionts or their associations can be strengthened from analyses of seasonal patterns and their comparison with expectations from laboratory experiments. In studies of seasonal dynamics, the effect of drift is ideally ruled out using spatiotemporal replication. For example, Smith *et al*. (2015) reported correlated change in the symbiont frequencies and the parasitoid-induced host mortality which, together with the laboratory evidence for symbiont-conferred resistance against parasitoids, suggested a causal relationship between them. Also, Montllor *et al*. (2002) reported an increase of the frequency of *S*. *symbiotica* correlated with temperature, which was consistent with this symbiont helping to tolerate heat stress. Our sampling design was not suited for such inference, but the result that *H*. *defensa* was more abundant in summer than in spring (Fig. S1) was at least consistent with selection by parasitoids as also reported by Smith *et al*. (2015). Field observations are also informative when they do not match the expectations from laboratory work. For example, laboratory experiments suggested that *X*-*Type* does not provide any detectable benefit to the pea aphid, but it is quite frequent and positively associated to *H*. *defensa*, suggesting it might have benefited from hitchhiking during the spread of *H*. *defensa* (Doremus & Oliver 2017). Also, Wulff *et al*. (2013) did not find that the symbiont *Arsenophonus* was protecting its *Aphis glycines* host against its main parasites, but it was present at high frequency. This discrepancy between observation and expectation motivated further experiments revealing that *Arsenophonus* provides a general – yet to be described – benefit to the aphid (Wulff & White 2015). Although difficult to interpret, field surveys remain crucial for our understanding of the ecology of symbioses.

## Conclusion

The fate of holobionts depends on host-symbiont interactions as well as on symbiont-symbiont interactions, but identifying them is not always straightforward. The approach consisting in analysing the frequency of associations in the field is useful. However, the results it yields must be interpreted carefully, in particular in the case of maternally transmitted symbionts, as patterns expected to be produced by interactions between symbionts are also induced by drift. This highlights the need for translating the biological null hypothesis into a valid statistical null hypothesis. Failure to do so may result in biological misinterpretations of survey results.

## Acknowledgements

We are very grateful to Paula Rodriguez for her help with insect rearing and to Marco Thali and Joelle Schmid for assistance with the molecular analyses of the field samples. We also thank Ailsa McLean for sharing several *Spiroplasma*-infected aphid lines. This work was supported by a Sinergia grant from the Swiss National Science Foundation (grant nr. CRSII3_154396 to CV).

## Data Accessibility

The DNA sequences used in this study are available in Genbank (accession numbers: MG288511 to MG288588). The main dataset and the Appendices S1 and S2 containing additional results on the spatiotemporal distribution of the symbionts and the annotated code of the simulation model are deposited in the Dryad Digital Repository: DOI WILL FOLLOW WHEN ACCEPTED OR ASKED FOR BY EDITORS.

## Author Contributions

HK, CH, CV and HMH performed the field sampling; HK, CH and HMH carried out the molecular analysis of the field samples; HMH was responsible for the data analysis and developed the model; HMH, HK, CH and CV wrote the paper.

